# Fine-scale Population Structure and Demographic History of Han Chinese Inferred from Haplotype Network of 111,000 Genomes

**DOI:** 10.1101/2020.07.03.166413

**Authors:** Ao Lan, Kang Kang, Senwei Tang, Xiaoli Wu, Lizhong Wang, Teng Li, Haoyi Weng, Junjie Deng, WeGene Research Team, Qiang Zheng, Xiaotian Yao, Gang Chen

**Author notes:** These authors contributed equally to this work (Ao Lan, Kang Kang, Senwei Tang, Xiaoli Wu). Correspondence: Xiaotian Yao & Dr. Gang Chen.

## Abstract

Han Chinese is the most populated ethnic group across the globe with a comprehensive substructure that resembles its cultural diversification. Studies have constructed the genetic polymorphism spectrum of Han Chinese, whereas high-resolution investigations are still missing to unveil its fine-scale substructure and trace the genetic imprints for its demographic history. Here we construct a haplotype network consisted of 111,000 genome-wide genotyped Han Chinese individuals from direct-to-consumer genetic testing and over 1.3 billion identity-by-descent (IBD) links. We observed a clear separation of the northern and southern Han Chinese and captured 5 subclusters and 17 sub-subclusters in haplotype network hierarchical clustering, corresponding to geography (especially mountain ranges), immigration waves, and clans with cultural-linguistic segregation. We inferred differentiated split histories and founder effects for population clans Cantonese, Hakka, and Minnan-Chaoshanese in southern China, and also unveiled more recent demographic events within the past few centuries, such as *Zou Xikou* and *Chuang Guandong*. The composition shifts of the native and current residents of four major metropolitans (Beijing, Shanghai, Guangzhou, and Shenzhen) imply a rapidly vanished genetic barrier between subpopulations. Our study yields a fine-scale population structure of Han Chinese and provides profound insights into the nation’s genetic and cultural-linguistic multiformity.

## INTRODUCTION

Population genomics has provided magnificent insights into the evolutionary pathway and the genetic composition of human beings. The prior large-scale studies, such as the 1000 Genomes Project (1KGP) (1000 Genomes Project Consortium et al., 2015), have predominantly centered on the variation spectrum in human genomes, which empowered the recognition of the genetic divergence of various populations across the globe. Comparing with the variation-scale profile, the haplotype sharing network within a population may administer a finer resolution for discriminating the substructures elicited by recent demographic events such as migration, admixture, segregation, and natural selection (Palamara et al., 2012; Powell et al., 2010; Speed and Balding, 2015). As two pilot studies, the geographical subpopulation structures of the British and Finnish populations have been well-demonstrated (Leslie et al., 2015; Martin et al., 2018). AncestryDNA, a direct-to-consumer genetic testing (DTC-GT) service provider, also published the fine-scale population structure in North America from their *in-house* biobank (Han et al., 2017).

As one of the most ancient nations, China is populated with the world’s largest ethnic group, Han Chinese. It is of great concern to conduct comprehensive genomics research to testify the nation’s historical records and legends, mine undocumented demographic events, and map its cultural diversification with the genetic imprints. Former microarray-based studies have identified an evident north-south genetic differentiation of Han Chinese (Chen et al., 2009; Xu et al., 2009). The low-coverage sequencing of over 11,000 Han Chinese uncovered a population structure along the east-west axis (Chiang et al., 2018). The deep sequencing of over 10,000 Chinese has provided extensive genetic markers of high quality (Cao et al., 2020). However, the limited sample volume of these studies remains insufficient for a highly modularized nationwide haplotype network, and the hospital-based cohort may also skew toward region-specific subpopulations. The largest published population study of the Chinese people has utilized the ultra-low depth sequencing data from the non-invasive prenatal testing to establish the nation-wide SNP spectrum (Liu et al., 2018), but lacks the resolution on an individual scale. Nevertheless, these datasets cannot simultaneously afford sufficient sample size, dense genetic markers to assemble shared haplotypes, and a well-proportioned participant distribution across the country to unscrew the subpopulation structure. A whole-genome genotyping dataset from a country-wide DTC-GT service provider is still an ideal solution to balance the cost of effect of haplotype network construction on a national scale.

In the present work, we create the haplotype network from the identity-by-descent (IBD) segments shared by 110,955 consented DTC-GT users from WeGene, China. We identify and annotate the subpopulation partitions using a hierarchical clustering approach and map the genetic separations with linguistic and cultural differentiation or historic demographic events.

## RESULTS

### Study Participants and the IBD Network Features

The 110,955 consented participants with self-reported ethnicity, birthplace (in prefecture-level), and current residence were recruited from the WeGene Biobank (**Figure S1**). All participants were genotyped with one of two custom arrays: Affymetrix WeGene V1 Array or Illumina WeGene V2 Array. After quality control, we utilized 350,140 autosomal single nucleotide polymorphisms (SNPs) to identify IBD segments (**Figure S2**). We then yielded a haplotype network composed of 102,822 vertices and 1.3 billion edges (total IBDs with a minimal length of 2 centiMorgan between a pair of individuals).

The principal component analysis (PCA) of the SNP profiles of the Han Chinese individuals resembles previous population studies (Cao et al., 2020; Liu et al., 2018), with similar proportions of variance explained by the first two PCs (0.25% and 0.13%) (**Figure 1a**). The PCA analysis of the IBD profiles exhibits a better separation among individuals from different geographical regions (**Figure 1b**). Also, higher proportions of variance were explained by the first two PCs of IBD (3.06% and 1.65%). IBD sharing indices were calculated between pairs of prefectures. The IBD-based genetic distance (IBD distance, calculated as 1 − IBD sharing index), SNP-based genetic distance (fixation index, *F*_*ST*_), and the geographical distance between cities highly correlate with each other (Pearson’s correlation, *p* < 0.01) (**Figure 1c**). As the *F*_*ST*_ distribution was heavily right-skewed and the IBD distance emerges while *F*_*ST*_ remains low (Pearson’s correlation between squared IBD distance and *F*_*ST*_: 0.81), the IBD dissimilarity has the potential to achieve a higher resolution among the communities with similar genetic backgrounds. Both SNP- and IBD-based genetic dissimilarity projections (**Figure 1d-e**) are associated with the cities’ spatial distribution (**Figure S3**), while the IBD analysis has presented better modularity for the prefectures from the same region (**Figure 1e**): for instance, the southern prefectures (yellow nodes) are immensely placed in specified modules in the IBD distance projection (ANOSIM test among the three southern provinces, R = 0.55, *p* = 0.001), while such partitions were less perceptible in the *F*_*ST*_ projection (**Figure 1d**), though also statistically significant (ANOSIM test, R = 0.28, *p* = 0.001). These city modules may preferably pronounce the genetic segregation among distinguished clans (Canton, Hakka, Min-Chaoshan, and Guangxi). Greater differentiation was also captured by the IBD distance between northern and northeastern China (ANOSIM test, R = 0.29 for IBD distance and 0.12 for *F*_*ST*_, *p* = 0.001).

**Figure 1.**
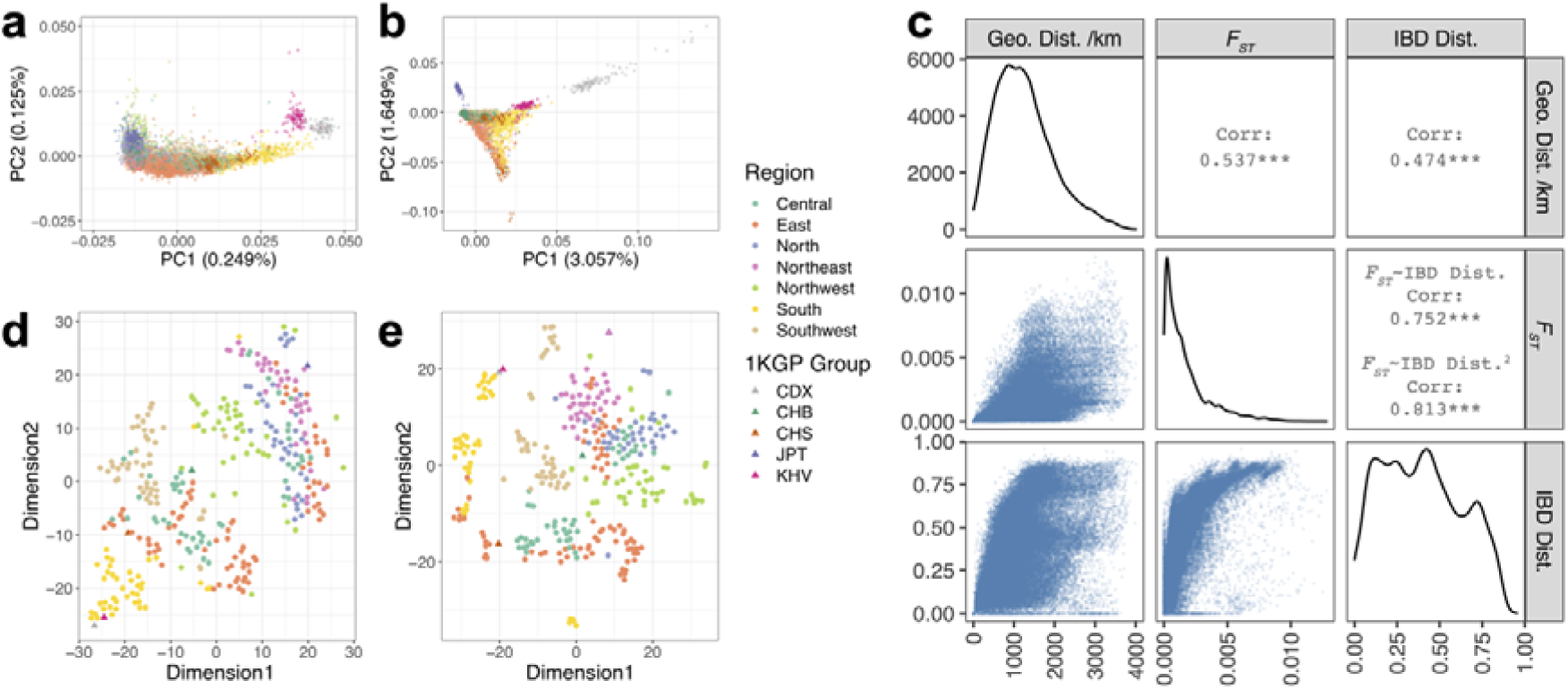
The genetic dissimilarities among individuals and among different cities in China. **a.** The PCA analysis of the SNP profiles of 5,000 randomly subsampled Han Chinese and 502 East Asian (EAS) samples from the 1000 Genomes Project (1KGP). **b**. The PCA analysis of the IBD profiles of the 5,502 individuals used in panel (**a**). **c.** The correlation between the inter-prefecture *F*_*ST*_, IBD-based genetic distance, and geographic distance. **d**. The t-distributed Stochastic Neighbor Embedding (t-SNE) projection of the SNP-based genetic distances (*F*_*ST*_) between prefecture pairs. **e.** The t-SNE projection of the IBD sharing indices between prefecture pairs. Panels **a, b, d**, and **e** share the same legend.

### Population Structure and Demographic Events

Hierarchical clustering was applied to the haplotype network to obtain a fine-scale population substructure recursively. The haplotype network clustering yielded two major clusters harboring 61.8% and 36.6% of the vertices in the entire network, successfully divided the population into the northern (1^st^-Northern China) and southern Chinese (2^nd^-Southern China). The most abundant cluster in each prefecture was colored distinctly in **Figure 2a**. The second stage clustering divided the southern population into three subclusters: 2^nd^-Southeast, 2^nd^-South, and 2^nd^-Southwest, and separated the Yangtze River Delta region (2^nd^-East) from the other northern population (2^nd^-North) (**Figure 2b**). In the third stage, more detailed partitions could be identified (**Figures 2c-f, S4, and S5**), where the imprints from ethnic fusion, recent movement, and linguistic-cultural division were able to be detected.

**Figure 2.**
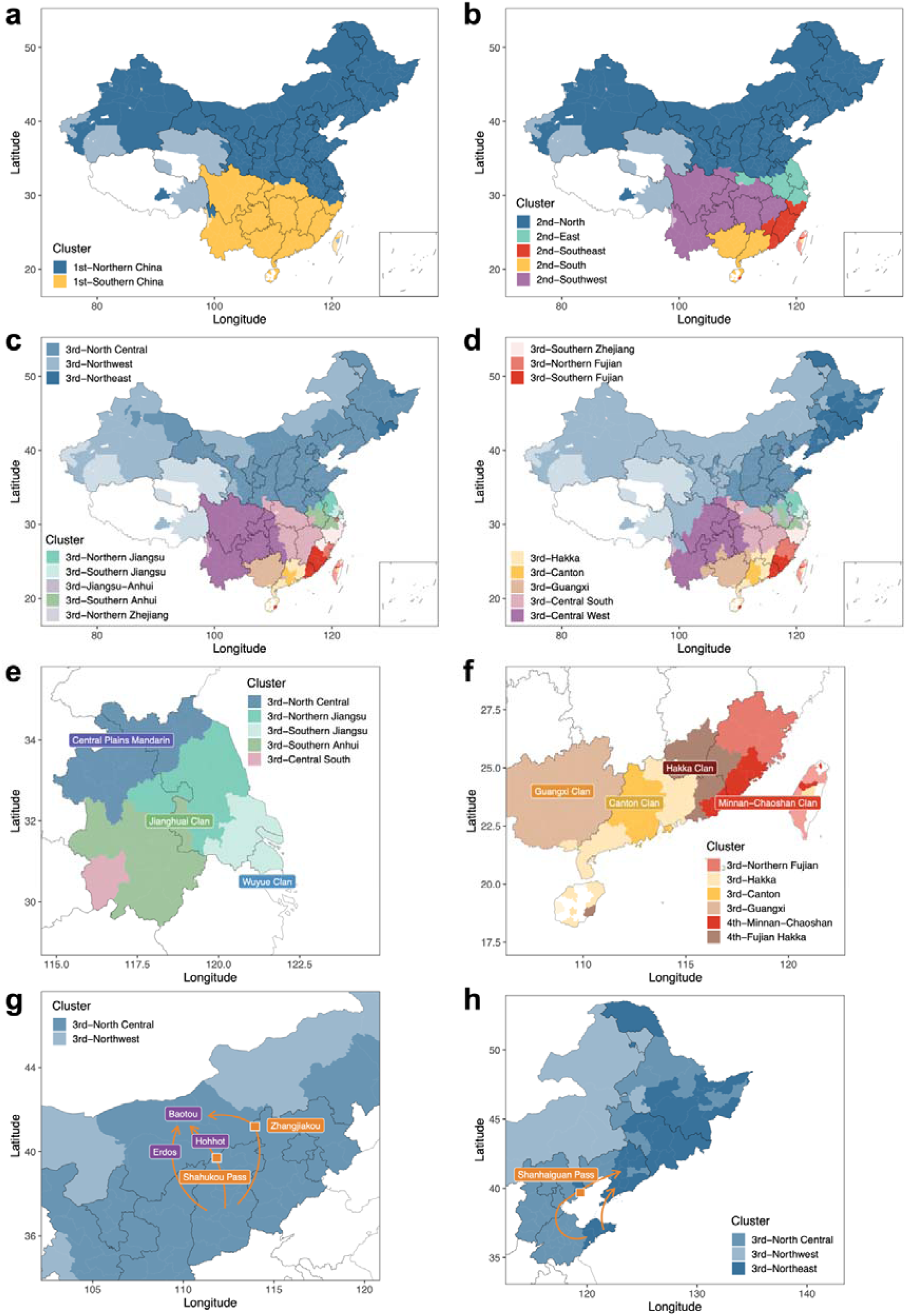
The hierarchical clustering of the haplotype network. **a-c.** The most populated 1^st^-level **(a)**, 2^nd^-level **(b)**, and 3^rd^-level **(c)** cluster in each prefecture. **d.** The 3^rd^-level cluster with the largest odds ratio in each prefecture. **e.** The spatial distribution of the 3^rd^-level subclusters in Jiangsu province is accompanying the linguistic-cultural division. The most populated clusters were shown. **f.** Three major clans in Guangdong, Canton, Hakka (falling into two clusters), and Min-Chaoshan, can be distinguished from the haplotype network clusters. The clusters with the largest odds ratio were shown. **g-h.** The paths of the *Zou Xikou* **(g)** and *Chuang Guandong* **(h)** migration waves. In panels **a-d**, 50% transparency was applied to the prefectures with small sample sizes (n < 10), and prefectures with no valid samples were left blank.

The subpopulation partitioning may attribute to an interplay of multiple factors including geography, politics, cultural, ethnic fusion, and natural selection. The southern boundary of the 2^nd^-North cluster is generally consistent with the Qinling-Huaihe Line (**Figure 2b**), the geographical dividing line for northern and southern China, as the two parts differ from each other in climate, staple crop and culture. Such separation was clearly pronounced by the haplotype cluster distribution in Jiangsu and Anhui provinces that locates in the Huai River basin, where the Wuyue clan, Jianghuai clan, and the central plain mandarin speaking regions could be distinguished (**Figure 2e**). In Guangdong province, the spatial division of the three major clans (Canton, Hakka, and Minnan-Chaoshan) could also be linked with distinct haplotype subclusters (**Figure 3f**). In the north, the pattern of the 3^rd^-Northwest cluster is substantially following the geographic placement of the Mongolic and Altaic ethnic minorities. The outlier in of the Hetao Plain in the central of Inner Mongolia, where the leading cluster assembles the Central Plains, may imply the historic migration wave *Zou Xikou* (go beyond the western pass) during the Qing dynasty (**Figure 2g**). Similarly, the Shandong Peninsula and most northeastern cities partook a common subcluster by the largest odds ratio, 3^rd^-Northeast (**Figure 2h**), which also implies the *Chuang Guandong* (rush beyond the Shanhaiguan Pass) immigration wave. In the PCA analysis for SNP of the individuals from the 3^rd^-North Central and 3^rd^-Northeast, no detachment could be discerned (**Figure S6**).

**Figure 3.**
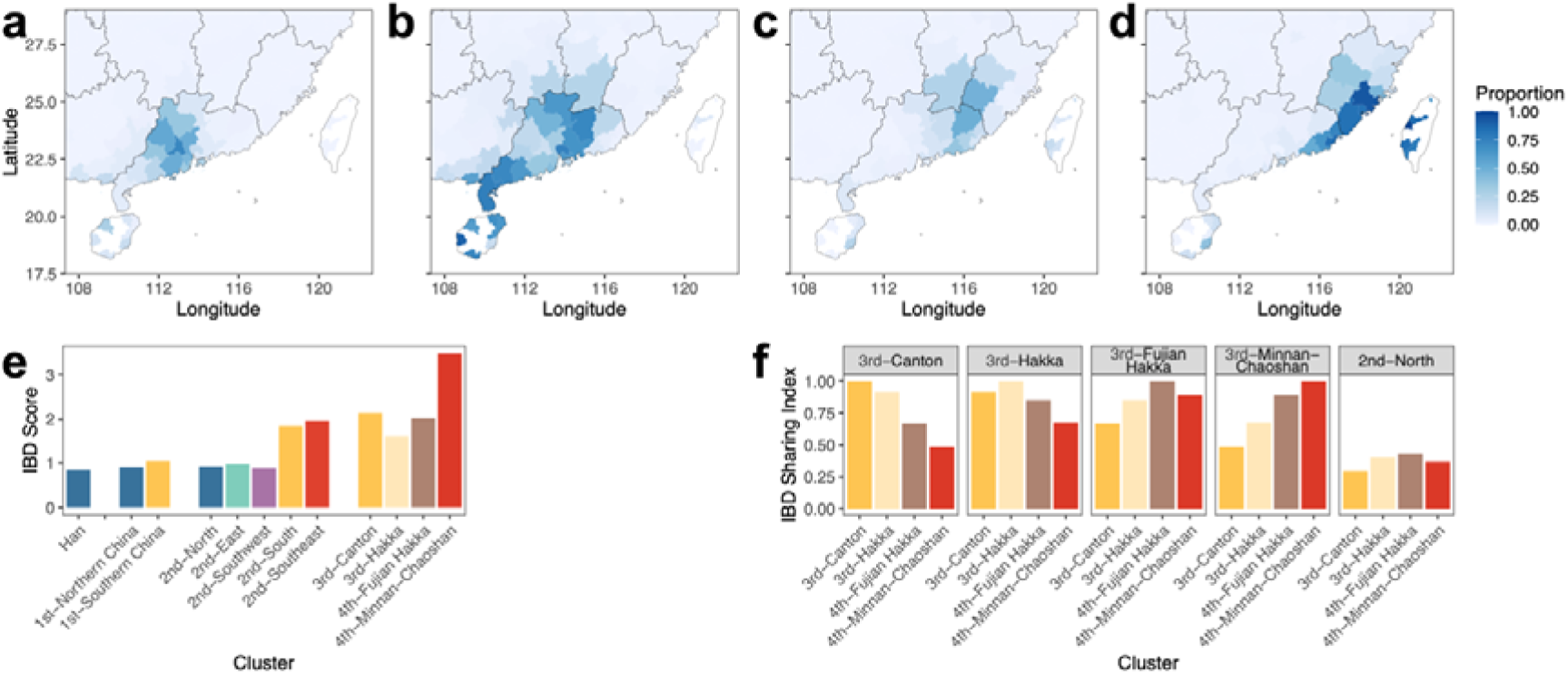
The distribution of the subclusters of the major clans in Guangdong and their population dynamics. **a-d.** Each subcluster’s population fraction in Guangdong province and adjacent regions. **e.** The IBD score of each subcluster. **f.** The IBD sharing indices between subclusters.

More subclusters could be classified in the south of the Qinling-Huaihe line (3^rd^-level subclusters, north: 3, south: 14). Guangdong and Fujian residents have formed various clans with specified languages, cultures, and habitations, and the differentiation is also portrayed by separate haplotype subclusters in this study (**Figure 3a-d**). Much higher IBD scores were observed in the 2^nd^-South (1.40) and 2^nd^-Southeast (1.57) populations (**Figure 3e**), particularly for the 4^th^-Minnan-Chaoshan subcluster (3.11), compared with the other clusters (for 2^nd^-North: 0.57, 2^nd^-East: 0.62, and 2^nd^-Southwest: 0.55, respectively). High IBD scores imply strong founder effects for these Han subpopulations, in line with the historic records for their southward migrations. In the meantime, the IBD sharing index between 3^rd^-Canton and 4^th^-Minnan-Chaoshan (0.41) was lower than the median IBD sharing index between two random clusters (0.61, one-sample Wilcoxon signed-rank test, p < 2 × 10^−16^) (**Figure 3f**), suggesting a high genetic disparity between these clans, though residing in adjacent regions for over a thousand year. The two Hakka subclusters exhibit the highest IBD sharing with the 2^nd^-North cluster (0.40 and 0.43), while 3^rd^-Canton shared the least (0.29).

### Modern Population Flows

In the contemporary era, economics is also shaping the new population substructures at an exceptionally rapid pace. We analyzed the modern population flows by comparing the participants’ birthplaces and current residences. In the four major metropolitans in China, most of the native residents (classified by participants’ birthplace) belong to the local cluster and subclusters (**Figure 4**): for instance, 86.8% of the Beijing native residents belong to the 2^nd^-North cluster; 51.5% of the Guangzhou native residents were members of 3^rd^-Canton and 30.8% were from 3^rd^-Hakka. However, the compositions of all these cities soon become an admixture of immigrants cross the country (**Figure 4**): only 17.5% and 21.3% of the current Guangzhou residents remain members of 3^rd^-Canton and 3^rd^-Hakka; the youngest one, Shenzhen, whose *de jure* population emerged from 0.3 million to over 20 million in the past 40 years, the fraction of the former dominant subcluster 3^rd^-Hakka reduced from 62.4% to only 13.3%. Accordingly, the 3^rd^-level cluster alpha-diversity (Shannon index) of these four cities increased from 1.47 ± 0.34 to 2.25 ± 0.25 (one-tailed, paired sample *t*-test, *p* = 0.003).

**Figure 4.**
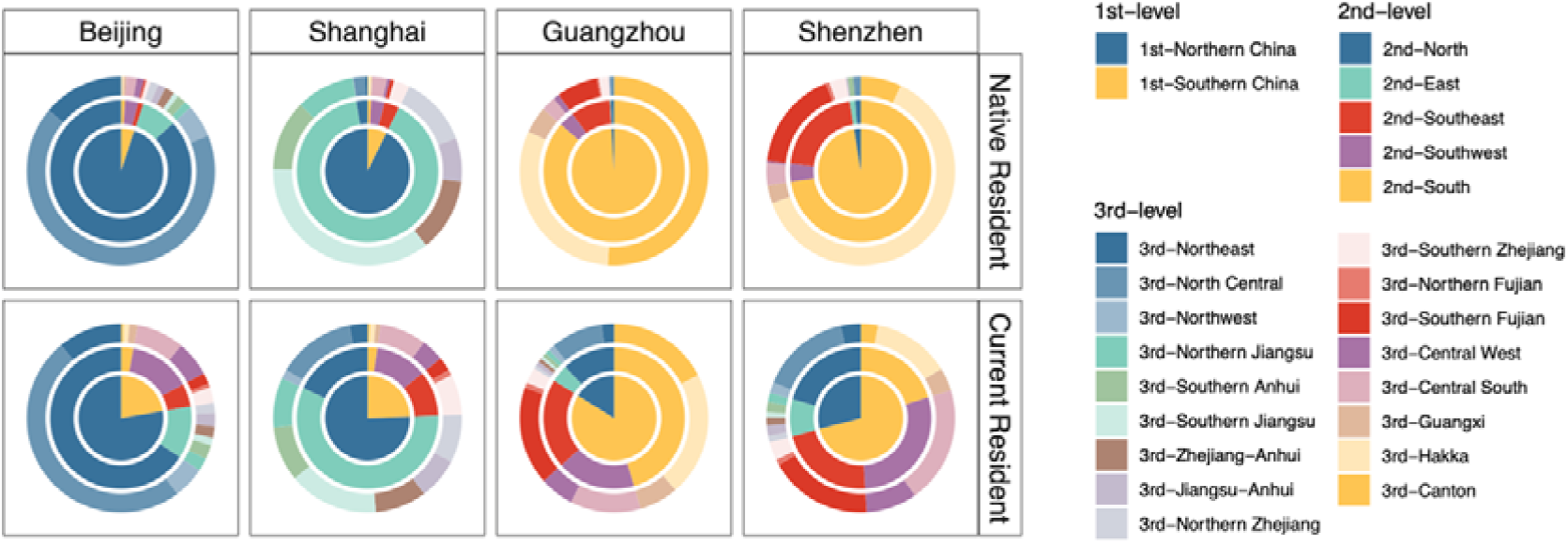
The resident composition by IBD network clusters in four major metropolitans, in comparison to the native residents (according to the birthplace) and the current residents. The inner to outer circles represent the compositions of the 1^st^, 2^nd^, and 3^rd^-level clusters respectively.

## DISCUSSION

As a biobank-scale population study of Chinese, we revealed the fine-scale subpopulation structure of Han Chinese by constructing a haplotype network of 110,000 genomes. The haplotype network shows a marked dependency between genetic distance and geography, but also process a step further to disclose the population substructures derived from recent demographic events or cultural and linguistic separation.

Previous large-scale studies on the Chinese population did not reach a fine-scale resolution for the population substructure, due to the limitation of sample size or genetic marker quantity and quality (Cao et al., 2020; Liu et al., 2018; Xu et al., 2009). The current biobanks in China also lack essential volume or nationwide representativeness of participants: only 6.3% of the 510,000 China Kadoorie Biobank participants were genome-wide genotyped (Chen et al., 2011), while the Taiwan Biobank project only sampled Han Chinese residing in Taiwan (Chen et al., 2016; Fan et al., 2008). Hence, the whole-genome genotyping dataset from DTC-GT services becomes a preferred solution to reveal the subpopulation structure that balanced the issues of participant distribution, sample size, genotyping cost, and marker density.

Unlike the North American haplotype network constructed from close relatives to reveal the post-Columbus population expansion (Han et al., 2017), we employed full-spectrum IBD pairs to trace the demographic events over a longer timescale. This enables founder effect estimation and cross-community dissimilarity analysis, which successfully revealed the genetic disparity among the clans in south China.

### Discrepant Application Scenarios between SNPs and IBDs

In the present study, the haplotype network and the SNP spectrum have provided related but independent information. In some scenarios, the SNP-based analysis lacks the essential resolution to subdivide population substructures with similar genetic makeup: for instance, the 3^rd^-Northeast clustering harboring the *Chuang Guandong* offsprings could not be distinguished from the other northern Chinese by SNPs.

### Geographical Impacts: Mountain Range > Climate > River

Mountain ranges have predominantly shaped the partition of the population substructure. Different subclusters with a considerable genetic distance reside on both sides of the major mountain ranges, such as the Qinling Mountains, Five Ridges, Wuyi Mountains, and Xuefeng Mountains. Different climate zones, the temperate zone, and the subtropical zone, also harbor different subpopulations, as revealed by the population composition of Anhui and Jiangsu provinces. There is no significant geographical isolation in this region, while different clans with disparate languages or dialects have formed, which also correlates with the rice farming and wheat (or millet before the Bronze Age) farming regions: wheat was cultivated in the Central Plains Mandarin speaking region, Wuyue relies on rice, while Jianghuai formed a cline. On the contrary, the isolation effect of great rivers, for instance, the Yangtze River, was not observed: the two sides of the Yangtze river always resemble each other in the subpopulation compositions, no matter in its upper-middle reaches, or in the delta region.

### War, Migration, and Politics: Keys to Population Split and National Fusion

War is a critical factor for ancient immigration, population split, and fusion. The southern Han Chinese clans are purported to be offerings of diverse southward movements from Qin to Song dynasties (Meacham, 1999; Wen et al., 2004). Cantonese was purported to be originated between Qin (221 to 206 BC) to Tang (618 to 907 AD) dynasties; Minnan-Chaoshan formed between Jin (266 to 420 AD) and Tang dynasties; the Hakka clan was composed of various southward movements between Tang and Qing (1612 to 1912 AD) dynasties, with a relatively short history and manifold origins. These histories were supported by the IBD network analysis, where Hakka has the lowest IBD score, but the highest IBD sharing index with northern clusters, suggesting a relatively late split with the Central Plains population. Cantonese and Minnan-Chaoshanese, though reside in adjacent regions, exhibited notable disparity, supporting the different origins. The 3^rd^-Canton cluster’s low IBD sharing index with the northern communities may also suggest its oldest split time, which is in line with historic records.

The haplotype network also successfully unveiled more recent demographic events driven by politics. *Zou Xikou* and *Chuang Guandong* were the largest recent migration waves of Han Chinese majorly happening within the past centuries, driven by politics. The population increase in the Central Plains imposed much pressure on the authorities. As a consequence, the Qing regime released the immigration ban for the Han people to reside beyond the Great Wall, the former reserved land of the ruling ethnic groups, Man and Mongol. As a result of the demic diffusion of Han Chinese, most of the northeastern Han people are offsprings of the *Chuang Guandong* wave. In our study, the genetic relationship between the Shandong Peninsula, the major origin of *Chuang Guandong*, and the northeastern Chinese was disclosed. As the most populated cluster (3^rd^-North Central) differs from the cluster with the largest OR (3^rd^-Northeast) in northeast China, the two clusters may imply the offsprings of migrants from different migration waves or choosing different routes: the inland residents using the land route via the Shanhaiguan Pass, or the coastal migrants using the sea route and landed on the Liaodong Peninsula (**Figure 2h**). Similarly, the *Zou Xikou* migrants from Shanxi province settled down to the traditional Mongolic regions including Baotou and Hohhot and became the largest local population now (**Figure 2g**).

### The Rapidly Vanished Population Boundaries

Though the Chinese populations have comprehensive substructures involving its long history and cultural pluralism, the genetic divergence between subpopulations may vanish over the coming decades, which may resemble the national fusion process that happened in Hispanic Latin America. Out analysis of the shifts of the metropolitans’ residents has confirmed the irreversible trend. Admittedly, the user distribution of a DTC-GT service could heavily skew toward youngsters and the current residents of the most developed regions and cities (**Figure S1**), particularly new economic migrants, which may result in an overestimation of the level of population mixing. The rapidly growing economy, coped with the emerging transportation capacity, has been speedily eliminating the genetic barriers between subpopulations. As the admixture increases, it might become more difficult to trace the demographic histories of a nation from either SNPs or IBDs. In this golden time for human population genomics, biobanking and biobank-scale studies are essential to mining the memories coded in our DNA.

## METHODS

### Study Design

#### Participants

All participants involved in this study were drawn from consenting WeGene customers. Participants with self-reported ethnicity, prefecture-level birthplace, and current residence were included (n = 110,955), and the demographic data were collected in April 2020. The East Asian samples (EAS) from the 1000 Genomes Project (1KGP) (n = 504) were integrated into the database. Duplicated genetic profiles from the same individual (n = 144) and profiles with relatedness up to the second-degree kinship (n = 8,493) were identified with *King* V2.2.1 (Manichaikul et al., 2010) with default parameters and excluded from analyses. Finally, 102,822 genetic profiles were acquired for analyses.

#### Ethnic approval and compliance

Informed consent for online research was obtained from all individual participants included in the study. The study was approved by the Ethical Committee of Shenzhen WeGene Clinical Laboratory. The study was conducted following the human and ethical research principles of The Ministry of Science and Technology of the People’s Republic of China (Regulation of the Administration of Human Genetic Resources, July 1, 2019).

#### DNA sampling and genotyping assay

Saliva samples for DNA extraction were collected processed following the previously published protocol (Kang et. al, in press). Samples were genotyped on one of two custom arrays: Affymetrix WeGene V1 Array (596,744 SNPs) by Affymetrix GeneTitan MC Instrument, and Illumina WeGene V2 Array (742,762 SNPs) by Illumina iScan System. A minimal genotyping call of 98.5% was required for a valid sample.

### Data Processing

#### Genetic marker quality control

Indels, heterosomal loci, and loci with more than two allelic states were removed from the genotyping data. For both arrays, SNP markers were filtered with *Plink* V1.9 (Purcell et al., 2007) with parameters “*--maf 0.001 --geno 0.05*” respectively. Only the intersection of the two arrays with identical allelic states was retained. To minimize the impact of the batch effect between the two arrays, for each biallelic SNP, a Chi-square test was performed among the three genotypes, and the *p*-values were Bonferroni corrected. SNPs with significant batch effect (*false discovery rate (FDR)* < 0.01) were eliminated. PCA analyses for the SNP sets before and after batch effect removal were illustrated in **Figure S7**. The density of the SNP markers used for IBD detection was shown in **Figure S8**.

#### 1KGP sample integration

The genotypes of the selected genetic markers of the 504 EAS samples were extracted with *VCFtools* V0.1.15 (Danecek et al., 2011). The genotypes of SNPs with inconsistent allelic states with the WeGene samples were set to a missing value. Then the genetic profiles of the 504 EAS samples were concatenated with the WeGene samples.

#### Genotype phasing

For the WeGene samples and 1KGP samples, we employed Eagle V2.3.5 (Loh et al., 2016) for a reference panel-free genotype phasing, using the default parameters.

#### IBD detection and merging

To minimize false-positive haplotype sharing, we identified the IBD segments (with a minimal length of 1 cM) with *Refined-IBD* (Browning and Browning, 2013) with default parameters. We then merged adjacent IBD segments with a gap less than 0.6 cM and no more than one genotype discordance in the gap region as one consecutive IBD segment, using the *merge*-*ibd-segments* function. In sum, 4,585 million IBD segments were yielded.

#### IBD segment quality control

We exclude the IBD segments with overlaps with any of the following regions annotated by the UCSC hg19 reference genome (http://genome.ucsc.edu/): centromeres, telomeres, acrocentric short chromosomal arms, heterochromatic regions, clones, and contigs identified in the “gaps” table.

For each SNP marker, the amount of IBD segments harboring it was summarized as the IBD coverage. 25% and 75% quantiles (Q1 and Q3) and the interquartile range (IQR) were calculated. The regions with an IBD coverage ≥ 75% Q3 + 1.5 × IQR were marked as IBD hotspots (**Figure S9 and Table S2**). IBD segments fell in or overlapped with such IBD hotspots were discarded.

#### Hierarchical clustering

The haplotype network was constructed with edges representing and weighted by the total shared IBD length (≥ 2 cM) between each pair of individuals. For the detection of population substructures recursively, we retained the edges corresponding to a total IBD ≥ 3 cM and applied the Louvain method for the hierarchical clustering (Blondel et al., 2008). The R package *igraph* was employed to apply. The clustering was performed for five levels. If a cluster or subcluster contained less than 50 nodes or was composed with < 1% nodes of its parent cluster, or was the only subcluster of its parent cluster, its next-level clustering stopped. In the 3^rd^ to 5^th^ levels, a cluster might be subdivided into fragmented and meaningless subclusters. To avoid this, we summarized the node counts in a subcluster × prefecture matrix, and pairwisely calculated the Spearman’s correlation between subclusters. The subclusters with pairwise correlation coefficients ≥ 0.8 were merged as one subcluster and would not be subdivided during the next-level clustering.

In each prefecture, the proportions and odds ratios (OR) of each cluster were calculated. The dominating clusters were named according to the cluster’s geographical distribution. The statistics of major clusters were summarized in **Table S1**. The geographical distributions of the clusters were shown in **Figures S5 and S6**.

### Statistics

#### IBD score, IBD sharing index, and genetic distances

IBD score was introduced to represent the mean total IBD length among all individual pairs within a community, following the previously published method (Consortium, 2019). IBD scores were calculated for prefectures, clusters, ethnic groups, and community subsets. For community *i* with a size of *n*_*i*_, *k* and *l* are an individual pair belonging to community *i*, the IBD score of community *i* was calculated as:

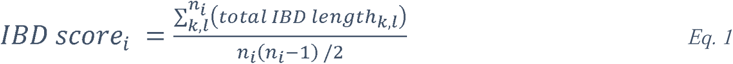

IBD sharing index was introduced to represent the mean total IBD length among all individual pairs from two communities and normalized by the IBD scores of the two communities to eliminate founder effects in different degrees. For community *i* and *j* with sizes of *n*_*i*_ and *n*_*j*_, respectively, *k* is a member of community *i* and *l* is a member of community *j*, the IBD sharing index between *j* and *j* was calculated as:

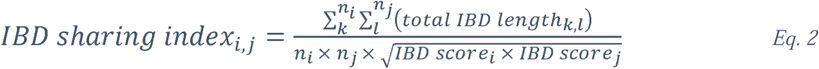

IBD distance between two communities was calculated as 1 − IBD sharing index. The IBD distances < 0 were rescaled to 0.

#### Data projection

Principal component analysis (PCA) was applied to the SNP profiles and the IBD profiles of 5,000 randomly subsampled Han Chinese individuals and the 502 EAS samples. For all quality-controlled SNPs, the redundant markers sharing the same linkage disequilibrium (LD) block were removed from the PCA analysis with *Plink* V1.9 (Purcell et al., 2007) with parameters “*--indep-pairwise 50 5 0.5*”. Finally, 138,725 SNP markers were retained for the PCA analysis for SNPs. For the IBD profiles, the IBD sharing matrix among the 5,502 individuals was used as the input. PCA analysis was performed with *GCTA* V1.9 (Yang et al., 2011) with the function *GCTA-PCA*.

The SNP-based inter-city genetic distance was calculated as the fixation index (*F*_*ST*_) using *VCFtools* V0.1.15 (Danecek et al., 2011). The SNPs used for *F*_*ST*_ calculation were the same SNP set for IBD detection. T-distributed stochastic neighbor embedding (t-SNE) was used for the genetic distances among cities.

#### Basic statistics and visualization

Data process, statistics, and visualization were performed using *R* and *R* packages including *igraph, vegan* (Oksanen et al., 2007), *reshape2* (Wickham, 2012), *tidyverse* (Wickham et al., 2019), *RCy3* (Gustavsen et al., 2019), *ggplot2* (Wickham, 2016), *ggally* (Schloerke et al., 2011), *ggtree* (Yu et al., 2017), *pheatmap* (Kolde and Kolde, 2015), *patchwork* (Pedersen, 2017), and *ggnewscale* (Campitelli, 2019).

#### Data availability

In light of our commitment to customer privacy and regulations from the Administration of Human Genetic Resource of China, we will not be publishing the raw data from WeGene customers. For the purpose of reproducing the analyses, we can share the haplotype network topology on request after a compliance review. For questions about the analyses in this research, please contact the WeGene Research Team by email.

## Acknowledgments

We thank all WeGene users who consented to share their genotype and demographic information for research purposes. We thank Prof. Dr. Chuan-Chao Wang from Xiamen University for his valuable suggestions and comments on this study. We also thank the employees of WeGene Inc. who contributed to the development of the infrastructure that made this research possible.

## Conflict of Interest

The authors AL, KK, ST, XW, LW, TL, HW, JD, QZ, XY, and GC work for WeGene (Shenzhen Zaozhidao Technology Co. Ltd. or Shenzhen WeGene Clinical Laboratory).

## SUPPLEMENTARY INFORMATION

This document includes 9 supplementary figures and 2 supplementary tables.

## Notes

### Summary of Updates

Corrected the digital gibberish in figures

