## Supplementary Figures & Tables for "Fine-scale Population Structure and Demographic History of Han Chinese Inferred from Haplotype Network of 111,000 Genomes"

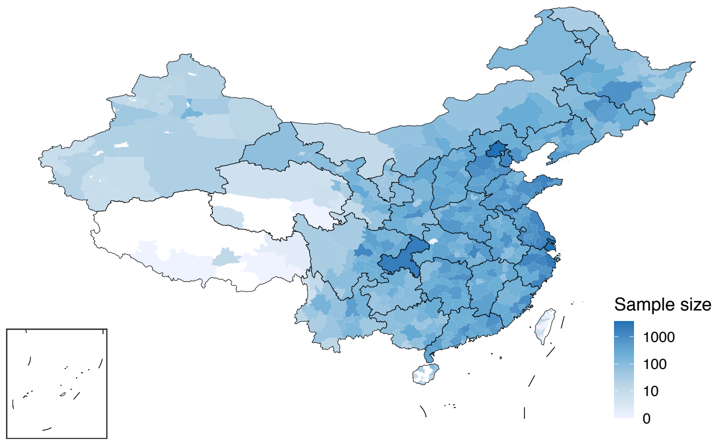


**Figure S1. The distribution of the participants' birthplaces.**

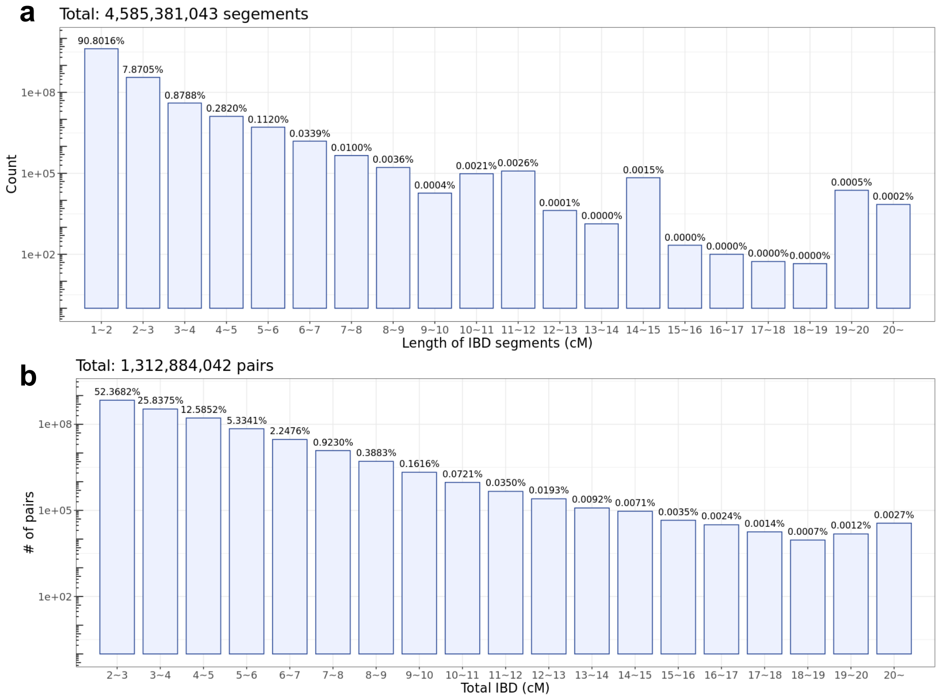


**Figure S2. The length distribution of IBD segments (a) and total IBD (b) shared by individual pairs.**

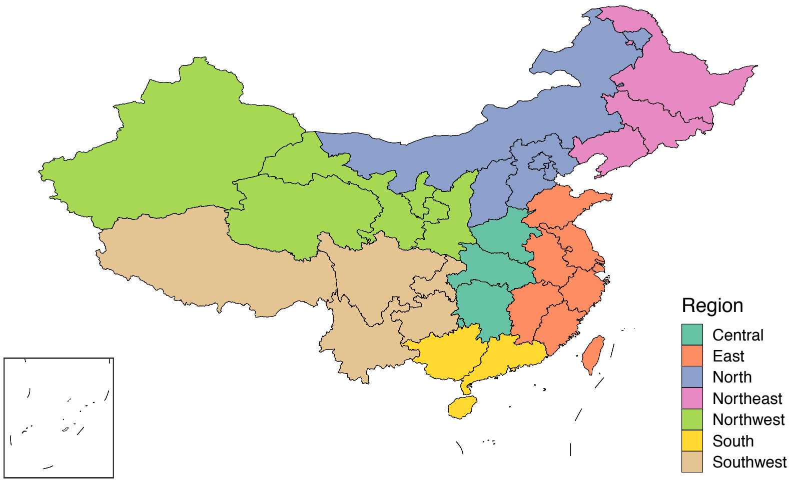


**Figure S3. The major geographic regions of China.**

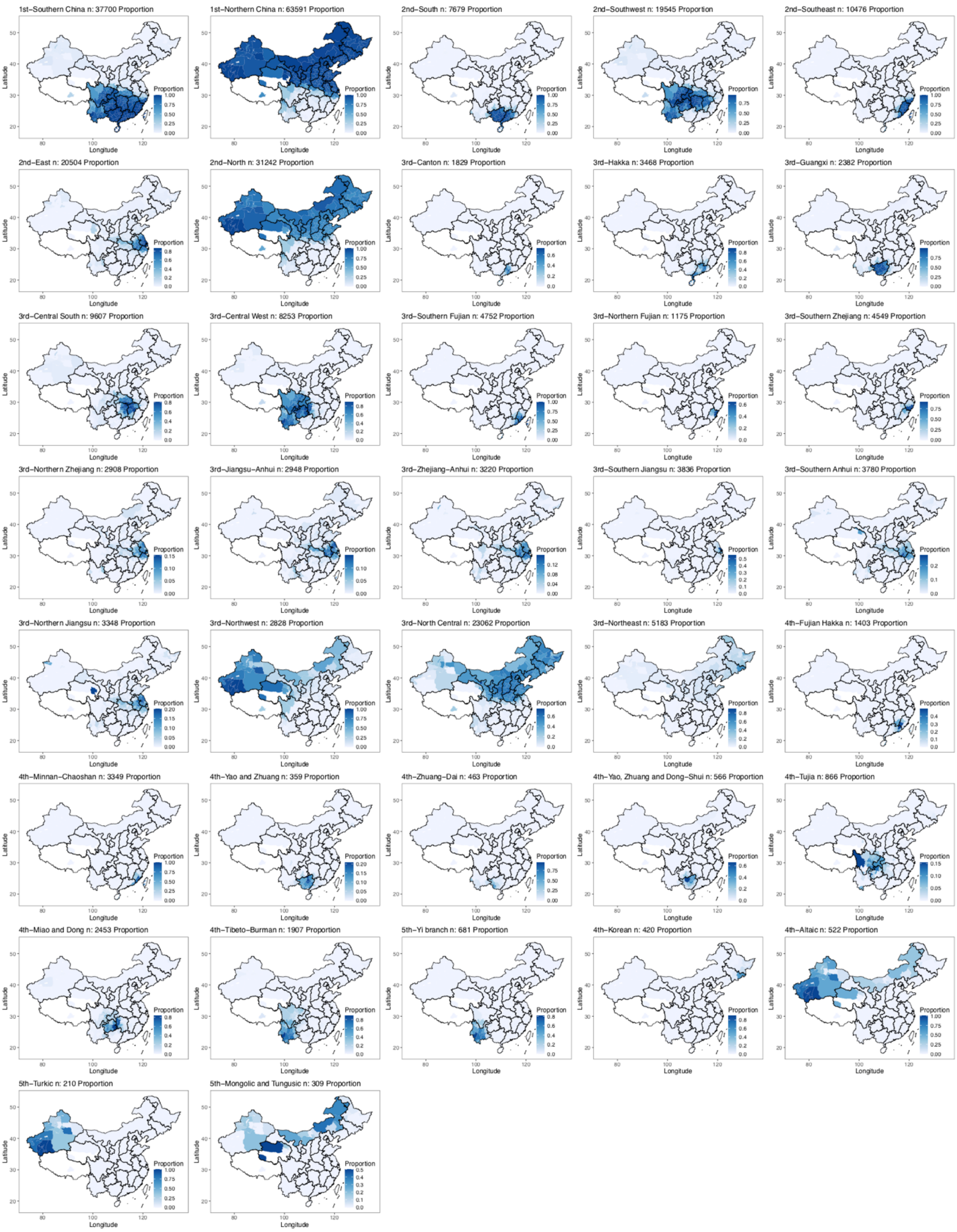


**Figure S4. The geographical distribution of the haplotype network clusters (local sample proportion by prefectures).**

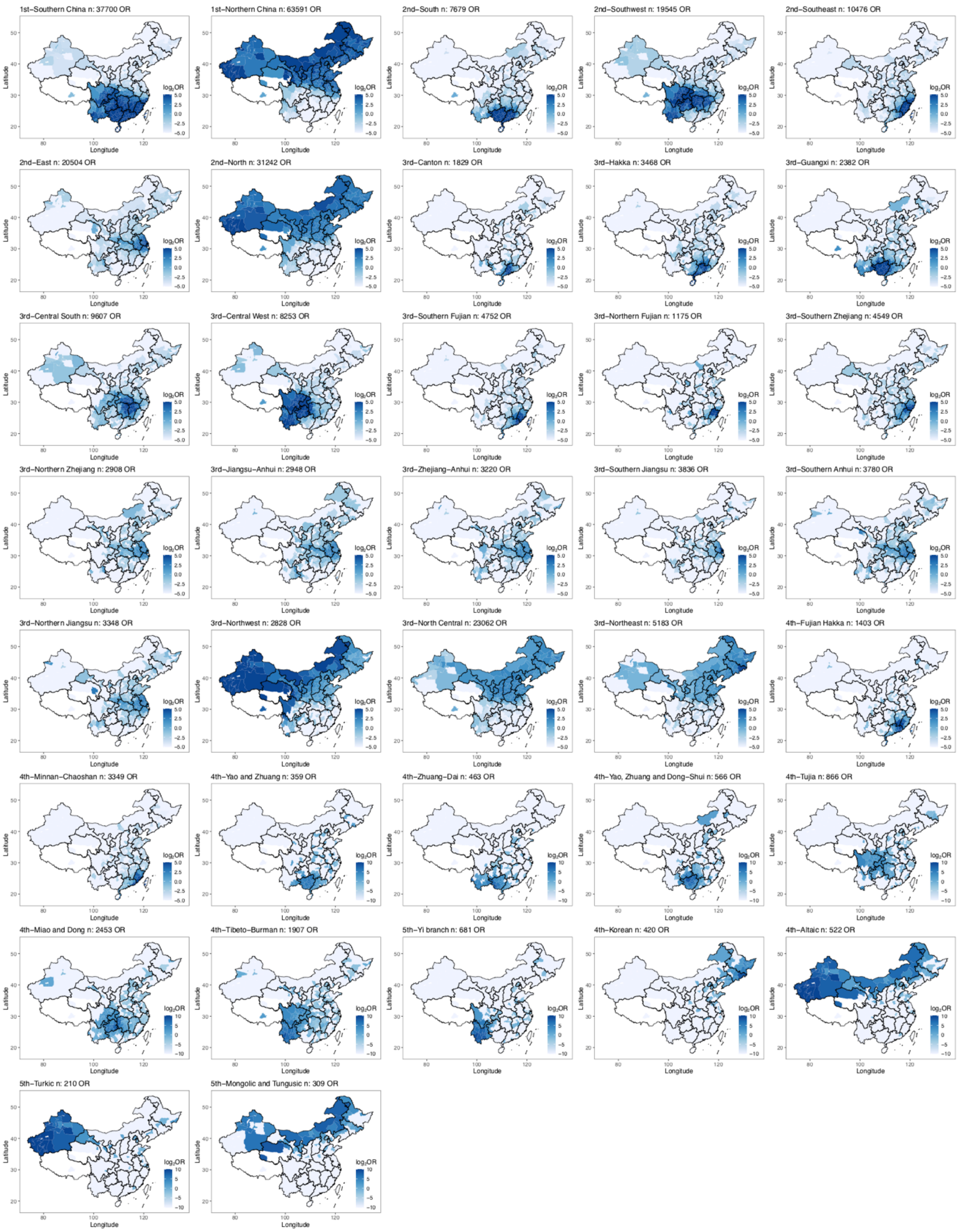


**Figure S5. The geographical distribution of the haplotype network clusters (odds ratios by prefectures).**

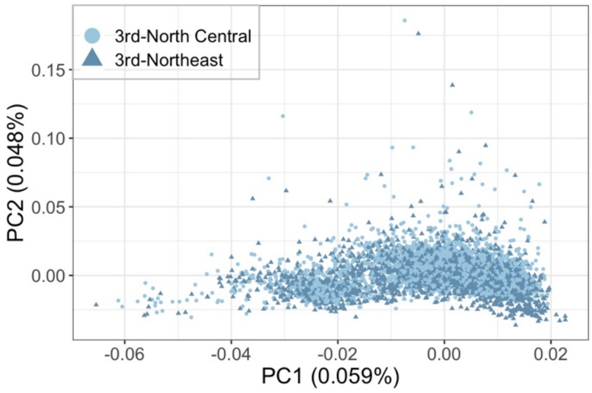


**Figure S6. The PCA projection of the SNP profiles of the members of 3rd-North Central and 3rd-Northeast.**

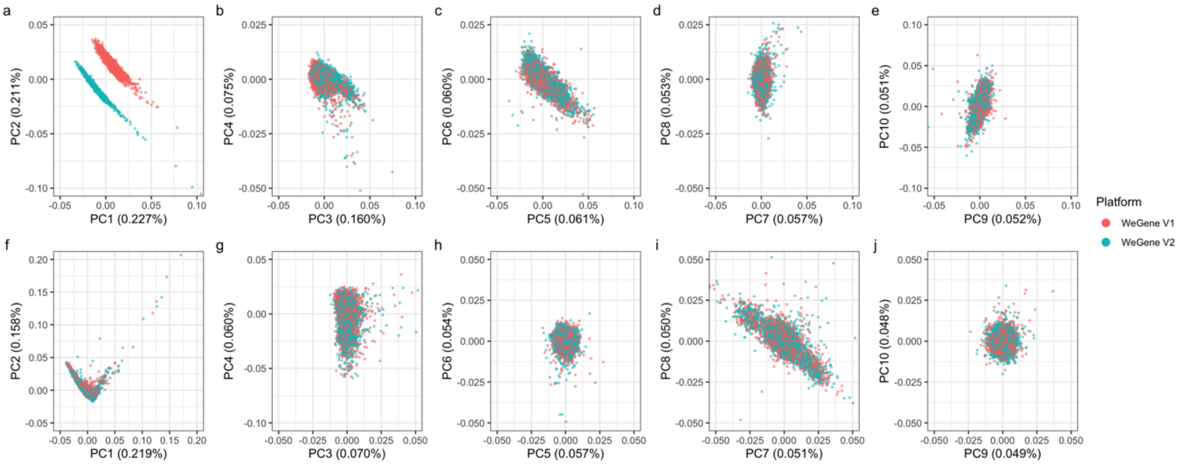


**Figure S7. PCA analysis of the SNP profiles from two arrays before (a-e) and after (f-j) the batch effect removal.** PCs 1 to 10 were shown for both the SNP sets before and after batch effect removal. The batch effect between the two microarrays could be observed in PCs 1 to 3 before the quality control (two-tailed *t*-test, $p < 1 \times 10^{-5}$), but the difference was efficiently eliminated after the quality control process (two-tailed *t*-test, $p > 0.05$). Outliers were excluded from visualization.

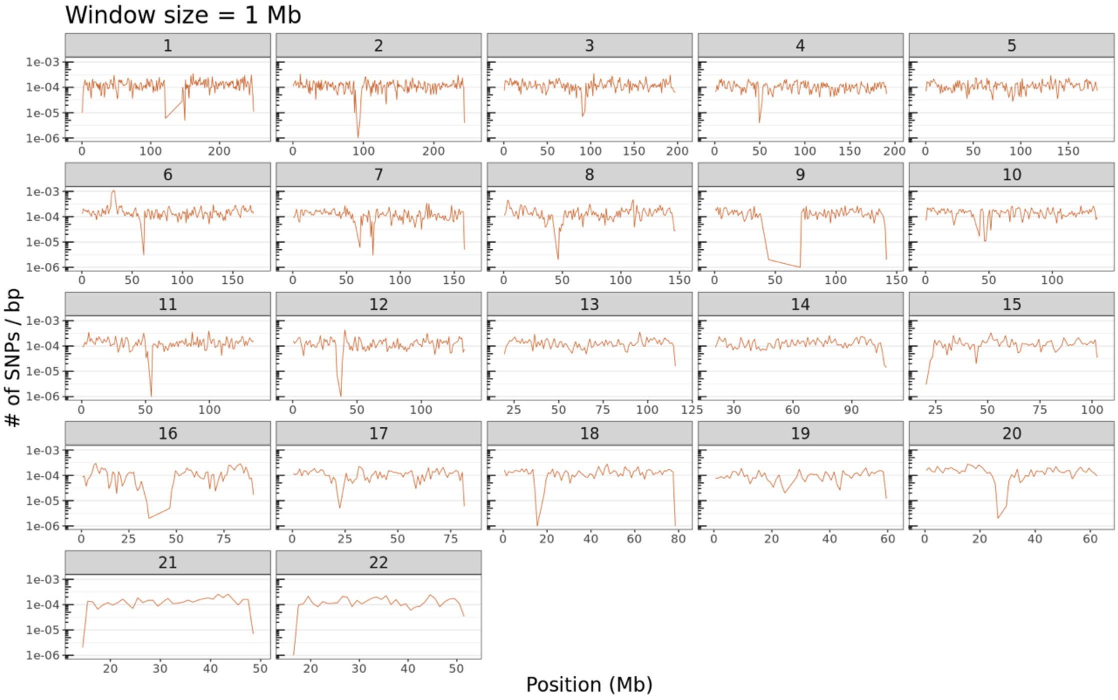


**Figure S8. The density of SNP markers selected for IBD detection.** SNP density was calculated for each 1 Mbp non-overlapped window.

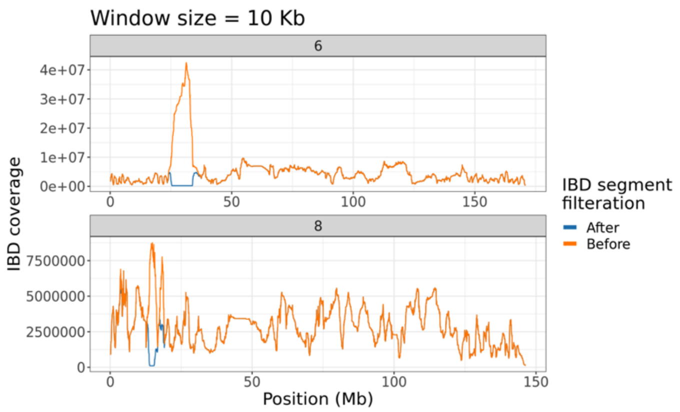


**Figure S9. An illustration of IBD hotspot removal.** Chromosomes 6 and 8 were illustrated as examples. The *HLA* region on Chr6 was an enriched region for IBD segments, thus removed from the down-stream analysis.

### SUPPLEMENTARY TABLES

**Table S1. The haplotype clusters.**

| Cluster Level and Name | # of Individuals |
|---|---|
| $1^{st}$-Northern China | 63591 |
| \|— $2^{nd}$-North | 31242 |
| \|— \|— $3^{rd}$-North Central | 23062 |
| \|— \|— $3^{rd}$-Northeast | 5183 |
| \|— \|— $3^{rd}$-Northwest | 2828 |
| \|— $2^{nd}$-East | 20504 |
| \|— \|— $3^{rd}$-Northern Zhejiang | 2908 |
| \|— \|— $3^{rd}$-Jiangsu-Anhui | 2948 |
| \|— \|— $3^{rd}$-Zhejiang-Anhui | 3220 |
| \|— \|— $3^{rd}$-Southern Jiangsu | 3836 |
| \|— \|— $3^{rd}$-Southern Anhui | 3780 |
| \|— \|— $3^{rd}$-Northern Jiangsu | 3348 |
| $1^{st}$-Southern China | 37700 |
| \|— $2^{nd}$-South | 7679 |
| \|— \|— $3^{rd}$-Canton | 1829 |
| \|— \|— $3^{rd}$-Hakka | 3468 |
| \|— \|— $3^{rd}$-Guangxi | 2382 |
| \|— $2^{nd}$-Southeast | 10476 |
| \|— \|— $3^{rd}$-Southern Fujian | 4752 |
| \|— \|— \|— $4^{th}$-Fujian Hakka | 1403 |
| \|— \|— \|— $4^{th}$-Minnan-Chaoshan | 3349 |
| \|— \|— $3^{rd}$-Northern Fujian | 1175 |
| \|— \|— $3^{rd}$-Southern Zhejiang | 4549 |
| \|— $2^{nd}$-Southwest | 19545 |
| \|— \|— $3^{rd}$-Central South | 9607 |
| \|— \|— $3^{rd}$-Central West | 8253 |

**Table S2. The IBD hotspots that removed from the analysis.**

| Chromosome | Start | End |
|---|---|---|
| **1** | 54,992,939 | 55,030,530 |
| **1** | 66,784,679 | 66,934,182 |
| **1** | 79,772,675 | 79,997,655 |
| **1** | 80,054,163 | 80,954,735 |
| **1** | 95,279,945 | 96,906,509 |
| **1** | 189,841,077 | 189,912,146 |
| **1** | 190,009,701 | 191,753,376 |
| **2** | 49,565,575 | 50,895,613 |
| **2** | 51,453,079 | 51,517,775 |
| **2** | 51,594,480 | 51,835,260 |
| **2** | 51,852,373 | 53,061,834 |

| | | |
|---|---|---|
| **2** | 53,123,426 | 53,212,803 |
| **2** | 53,263,553 | 53,358,953 |
| **2** | 57,397,237 | 57,465,539 |
| **2** | 57,665,702 | 59,356,000 |
| **2** | 166,421,414 | 167,506,029 |
| **2** | 168,412,798 | 168,479,374 |
| **2** | 205,851,076 | 205,889,646 |
| **2** | 205,906,945 | 205,912,403 |
| **2** | 205,917,744 | 206,345,419 |
| **3** | 81,962,522 | 82,249,474 |
| **3** | 82,279,967 | 82,293,860 |
| **3** | 82,441,812 | 86,526,422 |
| **3** | 86,531,505 | 86,537,262 |
| **3** | 86,554,319 | 86,752,414 |
| **3** | 87,090,544 | 87,110,674 |
| **3** | 87,134,800 | 87,173,324 |
| **3** | 87,219,009 | 94,867,122 |
| **3** | 94,983,900 | 95,028,989 |
| **3** | 95,122,880 | 95,163,558 |
| **3** | 98,869,960 | 99,320,707 |
| **3** | 99,383,411 | 99,391,933 |
| **3** | 128,988,045 | 129,298,994 |
| **3** | 129,300,291 | 130,512,274 |
| **4** | 97,568,242 | 101,260,798 |
| **5** | 111,911,276 | 112,004,034 |
| **5** | 137,754,695 | 138,719,526 |
| **5** | 160,388,599 | 160,412,665 |
| **6** | 25,411,464 | 33,854,816 |
| **7** | 9,998,779 | 10,721,233 |
| **7** | 21,351,951 | 21,496,397 |
| **7** | 21,657,572 | 21,682,636 |
| **7** | 21,694,832 | 21,745,879 |
| **7** | 52,748,225 | 53,239,490 |
| **7** | 57,395,834 | 57,401,925 |
| **7** | 62,705,018 | 63,566,551 |
| **7** | 63,628,981 | 66,482,088 |
| **8** | 3,699,398 | 3,706,150 |
| **8** | 13,905,760 | 15,469,237 |
| **8** | 15,479,638 | 15,486,397 |
| **8** | 15,656,128 | 15,765,369 |
| **8** | 18,228,116 | 18,444,535 |
| **8** | 18,460,837 | 18,535,156 |
| **8** | 18,668,681 | 18,674,816 |

| | | |
|---|---|---|
| **9** | 29,226,705 | 29,742,444 |
| **9** | 37,037,976 | 37,438,526 |
| **9** | 37,444,994 | 37,483,926 |
| **9** | 37,487,667 | 37,488,009 |
| **9** | 112,224,229 | 112,807,133 |
| **9** | 112,850,080 | 112,868,321 |
| **10** | 86,363,639 | 87,290,629 |
| **11** | 46,260,696 | 57,747,897 |
| **12** | 32,650,582 | 32,651,232 |
| **12** | 32,685,957 | 41,955,272 |
| **12** | 109,638,760 | 109,694,320 |
| **13** | 55,137,707 | 58,855,503 |
| **13** | 76,648,073 | 77,077,352 |
| **15** | 63,740,808 | 63,765,496 |
| **15** | 63,792,486 | 64,093,202 |
| **15** | 64,105,583 | 64,278,208 |
| **16** | 70,685,625 | 72,981,230 |
| **16** | 77,112,190 | 77,427,887 |
| **16** | 77,444,613 | 77,450,934 |
| **16** | 77,454,588 | 77,461,156 |
| **17** | 4,699,947 | 5,615,121 |
| **17** | 5,649,020 | 5,661,277 |
| **17** | 26,128,581 | 28,572,360 |
| **17** | 42,223,914 | 43,728,376 |
| **17** | 46,805,221 | 46,828,412 |
| **17** | 47,209,861 | 47,220,726 |
| **18** | 46,541,462 | 47,555,635 |
| **18** | 49,843,566 | 49,856,222 |
| **18** | 49,937,700 | 49,953,847 |
| **18** | 49,977,044 | 49,980,612 |
| **19** | 23,086,017 | 29,226,904 |
| **19** | 29,364,908 | 29,486,865 |
| **19** | 38,042,814 | 39,119,494 |
| **20** | 14,926,333 | 14,949,763 |
| **20** | 40,987,563 | 41,174,105 |
| **20** | 41,189,977 | 41,205,313 |
| **20** | 41,210,746 | 41,280,315 |
| **21** | 26,514,597 | 26,769,928 |
| **21** | 26,825,565 | 27,496,667 |
| **21** | 30,741,454 | 30,914,526 |
| **21** | 35,227,484 | 35,228,502 |
| **22** | 28,188,203 | 28,360,869 |
| **22** | 29,190,471 | 32,696,736 |

45
